# A run-length-compressed skiplist data structure for dynamic GBWTs supports time and space efficient pangenome operations over syncmers

**DOI:** 10.64898/2026.03.26.714584

**Authors:** Richard Durbin

## Abstract

Skiplists (Pugh, 1990) are probabilistic data structures over ordered lists supporting 𝒪 (log *N*) insertion and search, which share many properties with balanced binary trees. Previously we introduced the graph Burrows-Wheeler transform (GBWT) to support efficient search over pangenome path sets, but current implementations are static and cumbersome to build and use. Here we introduce a doubly-linked skiplist variant over run-length-compressed BWTs that supports 𝒪 (log *N*) rank and access operations, and a dynamic version of this that supports 𝒪 (log *N* + *S*) rank and insert operations, where S is the number of symbols in the alphabet. We use these to store and search over paths through a syncmer graph built from Edgar’s closed syncmers, equivalent to a sparse de Bruijn graph. Code is available in `rskip.[ch]` within the `syng` package at github.com/richarddurbin/syng. This builds a 5.8 GB lossless GBWT representation of 92 full human genomes, single-threaded in 52 minutes, on top of a 4GB 63bp syncmer set built in 37 minutes. Arbitrarily long maximal exact matches (MEMs) can then be found as seeds for sequence matches to the graph at a search rate of approximately 1Gbp per 10 seconds per thread.

## 1 Introduction

The idea behind pangenomes is that, when analysing data from a new individual, we should make use of the genetic variation known to be present in a species or population, rather than use a fixed linear reference genome sequence [24, 2]. To do this we combine information from many independently assembled genomes, removing redundancy by collapsing together shared segments of sequence. A standard way to do this is in a graph, with shared sequence segments represented by vertices, which are connected by edges that indicate what can follow what. Both the vertices and the edges are bi-directional, like highways or rail lines, which can be travelled in one direction or the other, with the sequence for a vertex being reverse-complemented when traversed in the opposite direction.

Most pangenome graph applications assign equal weight to all routes across vertices that follow edges of the graph, or perhaps weight paths according to the frequencies of their vertices and/or edges in the set of reference genomes from which the graph was constructed. However there is longer range information in the reference genomes, based on shared haplotypes, that we want to use. In the railway analogy, the graph is the network, but to get from A to B you need to take a series of trains that follow established paths through the network. This longer range haplotype information is what enables powerful genotype imputation methods that have provided core computational infrastructure underlying genetic association studies on millions of samples over the last fifteen years[11].

Recent methods for genotype imputation[21] have exploited the positional Burrows-Wheeler transform (PBWT)[3], which provides efficient storage of and matching to haplotype sequence panels. Our goal is to support a similarly efficient and powerful framework for pangenome graphs, where sequences do not just vary at colinearly aligned binary variants. As a step in this direction, I introduce here a computational framework for efficient calculation over graph Burrows-Wheeler transforms (GBWTs), the corresponding data structure to the PBWT for pangenome graphs, and present a practical implementation.

Below I present first the use of graph Burrows-Wheeler transforms (GBWTs) to store and search paths through pangenome graphs, and then the use of a variant form of Pugh’s skip list[17], which I call Rskip, for efficient computation over pangenome GBWTs. Then I describe key features of my `rskip` implementation and its use in the `syng` package, which builds pangenome graphs in which the vertices represent overlapping *k*-mers selected using Edgar’s closed syncmer criterion[4]. This is followed by the results of applying `syng` to the Human Pangenome Reference Consortium phase 1 release of 92 full human genomes[10].

## 2 Theory

Here we discuss using GBWTs to represent a set of paths on a pangenome graph and provide efficient search over these. We will use the term ‘vertex’ for the elements of the graph that are associated with sequences, in our case syncmers, and ‘node’ for nodes in the internal Rskip data structure that we use to manage the routing tables for the paths, which happens to also be a form of graph.

### 2.1 Path traversal and search in a sparse graph via local GBWTs

Given a graph vertex with a set of paths running through it, and an ordering on each of the bundles of paths coming into the vertex from other vertices, it is natural to also list the vertices from which the paths come, and hence derive a global ordering of incoming paths (left side of Figure 1). Then a simple routing table of corresponding output nodes, as highlighted in purple in Figure 1), is all that is needed to specify the output vertex for each incoming path. Given that the paths within each outgoing bundle are ordered colinearly with incoming paths, as for the X’s and Y’s in Figure 1, adjacent paths in a bundle will share recent sequence, and by construction the routing table will correspond to the subsection of the Burrows-Wheeler (BWT) transform over sequences of vertices where the last symbol corresponded to the current vertex. This graph vertex BWT is called the Graph Burrows-Wheeler transform[22, 6].

**Fig. 1.**
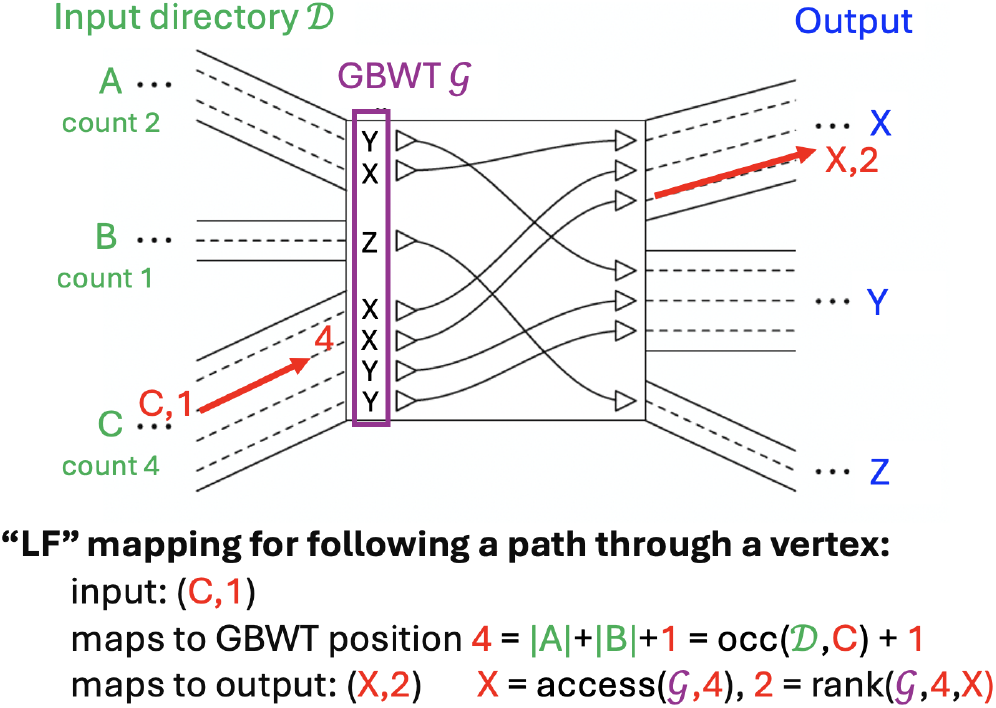
LF mapping to follow a path entering as offset 1 in the path bundle from C and exiting as offset 2 in the path bundle to X. The local GBWT 𝒢 provides the routing information, with the index in 𝒢 being the sum of the incoming offset and the total count of preceding symbols in the input directory 𝒟. All offsets and indexes are 0-based. Adapted from [22] with permission.

Because sharing past sequence is predictive of sharing future sequence and specifically the next symbol, BWTs in general and the GBWT in particular tend to contain runs of the same symbol. They are therefore efficiently compressed by run-length compression. The run-length compressed GBWT or rGBWT for Figure 1 would be `((Y,1),(X,1),(Z,1),(X,2),(Y,2))`.

When following a path using a GBWT we need operations `occ(`𝒟, *s*) on the input directory 𝒟 to give the total count of input paths from symbols listed before *s*, `access(`𝒢, *i*) to give the symbol at position *i* in the GBWT 𝒢 and `rank(`𝒢, *i, s*) to give the number of times *s* appears before position *i* in 𝒢, as shown in Figure 1. The same operations support efficient matching. There are standard algorithms linear in match length in terms of `occ()` and `rank()` operations to find the number of paths matching a substring or to find maximal exact matches (MEMs) of a longer search string (this is a little imprecise for MEMs, but let’s ignore that here).

Note that this construction shows that for the update operations we do not require counts for all vertex symbols, only the ones incident to the current vertex, nor a global ordering of the symbols, just a local ordering of those that are incident. This is helpful to us because there are many millions of vertices, and different orderings may be preferable for different vertices.

Typically BWTs are built statically by one process, and then an index structure is built over them by a different process to support the required operations. Here we introduce a novel data structure that naturally supports dynamic building of run-length compressed BWTs for efficient storage, traversal and search.

### 2.2 Skip lists for dynamic run-length-compressed arrays

If we don’t want to move large memory blocks when we insert into a dynamic array the simplest structure would be a linked list, but this incurs linear time traversal cost for all `access()` and `rank()`. There are many succinct data structures that can reduce these to 𝒪 (log *R*) (where *R* is the number of runs) or for some operations even constant time[15], but typically these assume relatively small alphabets, whereas our alphabet size is potentially unbounded, reaching in practice at least tens of thousands due to the repetitive structure of the genome (see Results).

Here we take advantage of the elegant skip list data structure introduced by Pugh in 1990[17], which augments a linked list with additional layers generated by a random process in which each node has an upward parent with probability *p* (Figure 2A). Skip lists provide a lightweight alternative to balanced trees, supporting `access()` in expected 𝒪 (log *R*) time as in Figure 2B, and also 𝒪 (log *R*) `insert()` (and if wanted `delete()`) by keeping track of the last node visited at each level on a stack and updating their pointers and counts as necessary for a new column. Storing run lengths is natural in a skip list because the nodes at upper levels already store counts.

**Fig. 2.**
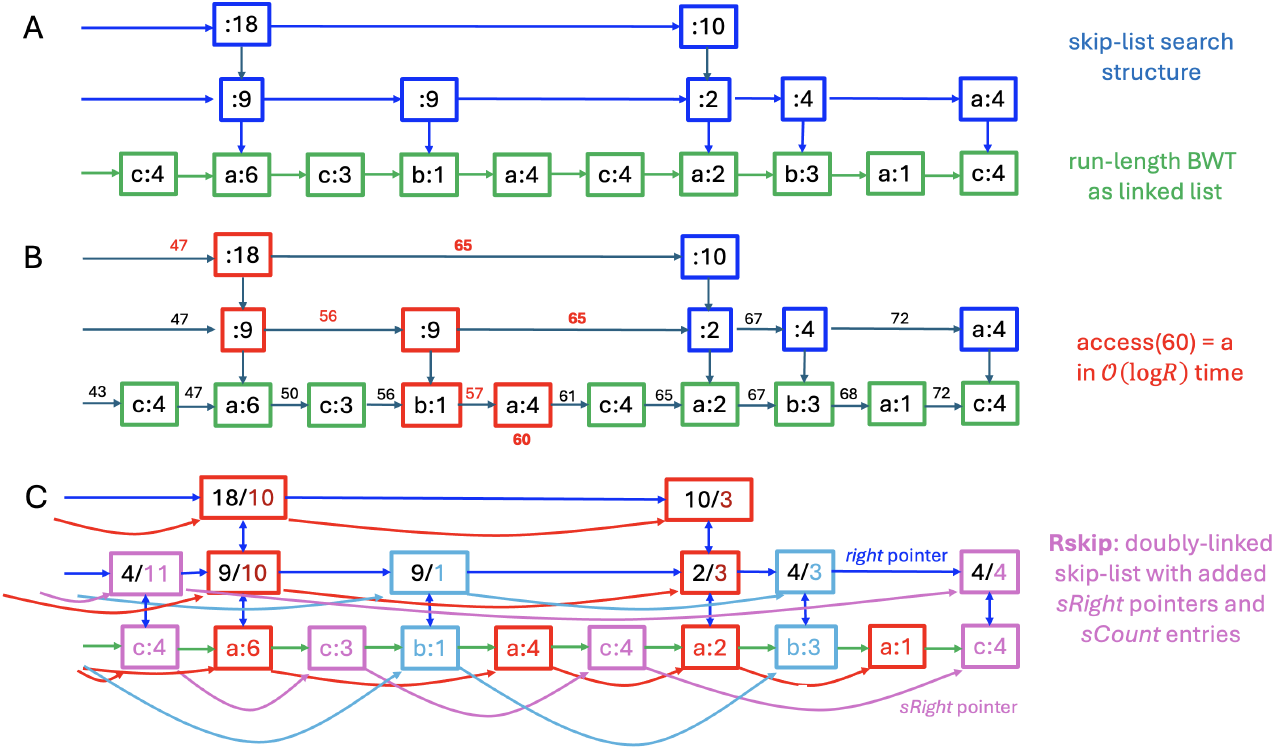
Skip list and Rskip data structures. A: a standard skip list over a run-length array, with columns up to (in this case) two levels. B: the `access()` operation is performed in expected 𝒪 (log *R*) time by starting at the top of the first column, moving right accumulating a partial sum, and dropping levels when the sum for the next right edge would exceed the target; only the partial sums coloured in red need to be calculated. C: Rskip structure with additional pointers and counts between nodes for the same symbol.

To support `rank()` once we are located at a node, we need to find the sum of counts of the same symbol in preceding nodes. We could count forward along the list how many are in succeeding nodes, then subtract from a total count that we maintain for each symbol, but this would be a linear time operation. Instead I add a second set of `right` pointers that point to the next node at the same level with the same symbol, accompanied by counts for how many of that symbol are present up to that node. Effectively this is embedding a skip list for each symbol within the same set of columns as in the original primary skip list. We also need to add `up` pointers in reverse to the `down` pointers. See Figure 2C. Now we can find the number of copies of symbol *s* by moving up and right along the `up` and `sRight` pointers, going up whenever possible, and accumulating `sCount`s as we go. This is exactly reciprocal to the initial `access()` search, so has expected time 𝒪 (log *R*_*s*_) where there are *R*_*s*_ runs for symbol *s*. Analogous to the `insert()` operation for a standard skip list, it is possible to also insert new columns into the Rskip structure: once the rank is known the relevant node is found again using `sAcccess()` on the symbol’s own skip list, and the stack of nodes up to that point can have their `sRight` pointers and `sCount` values updated.

Although the previous paragraph satisfies the requirement of `rank()` for LF mapping to traverse a path, where we start from a node with the required symbol, for general `rank(s,i)` for an arbitrary symbol *s* we would need to find the next node with that symbol, which will require a linear search on the bottom level. The expected time for this linear search given a random next symbol with the same frequency as in the reference is 𝒪 (*S*), where *S* is the alphabet size, since if symbol *s* has frequency *f*_*s*_ then the expected time for this search is 1*/f*_*s*_, so the expected time for a random symbol is ∑_*s*_ *f*_*s*_1*/f*_*s*_ = ∑_*s*_ 1 = *S*. Although this does not happen when traversing a path, it can happen when matching a new sequence, or inserting a new column. Fortunately, although this argument applies to a random input, and adversarial sequences could have much worse behaviour, empirically with real genome sequences the mean length searched is less than *S*. First, entries with the same symbol tend to be clustered - indeed this is what empowers run-length compression - and as a consequence the `rank()` operation is already located at the correct symbol, requiring no linear search.

Second, even conditional on needing to search, the mean linear search length is lower than *S*. I provide empirical evidence for these assertions in the Results section below.

In practice, I use two variants of the canonical Rskip structure described above. When building and requiring support for `insert()` I add back pointers `left` and `sLeft` to simplify the update code at the cost of some additional memory. In particular, when doing this there is no need to maintain a special start column for each symbol. For static use while searching, it is possible to replace the `count` and `sCount` in the nodes in Figure 2C by partial sums up to that node, in a single linear sweep through the structure. Then the `rank()` operation that follows `access()` during path following is a simple lookup, since the partial `sSum` of the symbol of a node is stored within the node. Furthermore, general `rank()` can be executed in 𝒪 (log *R*_*s*_) time by following the `sRight` and `down` pointers while looking at the `sum` entries. There is no longer an expected 𝒪 (*S*) term.

## 3 Implementation

The Rskip data structure and algorithms were implemented in C file `rskip.c` with public interface in `rskip.h`. These are available within the syng package at github.com/richarddurbin/syng, which implements pangenome construction and, for now, basic search operations based on rskip and Edgar’s closed syncmers[4].

### 3.1 Rskip implementation

To avoid memory fragmentation and ensure cache locality the entire Rskip data structure is held in a single contiguous memory block made of an array of `max` nodes. In static search mode I use five 32-bit integers per node of type `Fixed` to record `right, sRight` and `down` pointers as offsets into the array, and `sum` and `sSum` partial sums. On the bottom level we set a top bit of `sSum` as a flag and use the `down` slot to hold the symbol identifier `sym`. The first node of the block is used as a header, recording `max`, the number of symbols `nSym` and the starting node. This is followed by `nSym` nodes forming a directory for the symbols, which hold external symbol information and the total count for the symbol in the whole array. The next `nSym` nodes following the directory are the tops of the first columns for each symbol. Column height for new nodes is sampled from a geometric distribution with mean height 1.6 by iterative sampling from a lightweight random number generator so long as the returned value is under 3*/*8, which is close to the optimum 1*/e*.

The dynamic Rskip variant uses eleven integers per node: six for bidirectional pointers `right, left, sRight, sLeft, down, up`, four for counts before and after `before, count, sBefore, sCount` and one for `sym`. The `before/sBefore` counts are redundant with `[left].count/[sLeft].sCount` but simplify code at the left boundary which would otherwise require additional storage. There is again a header node and directory, in this case with the directory entries holding a pointer to the top of the left-most highest column for the relevant symbol. To manage new node allocation, a `free` pointer is kept in the header, and array size is doubled when more space is required

Because the Rskip structures are relatively heavy, I use a much simplified `Linear` node array for small run lists, in both static and dynamic modes. Specifically these contain a maximum of 128 two-byte nodes, with nodes holding either a one-byte symbol identifier and count, or if the count is 255 then the subsequent node holds a two-byte count. The header is now two nodes, and the directory uses four nodes per symbol. Again there is a `free` pointer in the header in dynamic mode.

Implicit in the description above is that internal to the Rskip I use symbol values denoted `sym` from the range `0..nSym-1`. Calls to the `rskip` package are made with a general integer, or in the case of `syng` with a general (symbol,offset) pair, and the corresponding `sym` is looked up/stored in the directory. Currently this lookup is done with a linear search, which in principle makes the time complexity 𝒪 (*S*) where *S* is the local alphabet size `nSym`. It would be possible to use a hash table or equivalent to make this constant time, but in practice the frequencies of symbols in pangenome graphs are very skewed (most genetic variants are rare). As a consequence, since the directory is built progressively as paths are added, high frequency symbols are at or close to the start of the directory list, meaning that the expected time for linear searches is short. Actual expected search times for a human pangenome are given in the results section.

### 3.2 syng implementation

Since the focus of this paper is on the Rskip data structure and algorithms, only brief details of the syng implementation are given here.

Syng pangenome graphs are bi-directional, with each vertex representing a syncmer and its reverse complement. Syncmers are fixed length *k*-mers with the property that one of their terminal *s*mers has less than or equal hash value compared to all internal *s*mers[4]. This gives them a guaranteed window property, like minimizers to which they are related[20], with consecutive syncmers in a sequence overlapping by at least *s* bases, though unlike minimizers syncmers are not context dependent: whether a *k*-mer is a syncmer is an intrinsic property of the *k*-mer sequence. Hence syncmers cover any sequence (except for its ends), and a list of syncmers together with offsets between them, plus terminal ends of length <= *k* − *s*, are sufficient to reconstruct the sequence. For syng we require that *k* is odd (default *k* = 63, *s* = 8) and use a positive syncmer index value to indicate traversal in the direction that lexicographically comes before its reverse complement, and minus that value for the reverse-complement direction. Syng interconverts sequences and syncmer lists via a rapid scanning algorithm and a hash table.

For each vertex syng keeps a 16-byte data structure with two sides, `in` and `out`, together with an 8-bit set of flags. In many cases there is only one edge for a side, termed a *simple* side; then the next syncmer index is stored directly in the side object, together with the offset to it and the edge count. Otherwise, a pointer to an Rskip is stored, which maintains the list of edges in its directory together with a BWT supporting path reconstruction.

To follow a path using LF-mapping with a BWT standardly requires a rank operation on the BWT and a sorted occurrence array. In our GBWT case only a small subset of the large total alphabet is relevant to each step, corresponding to the incoming and outgoing edges for each vertex in what is overall a very sparse graph. The rank operation is provided by `rsRank()` on the Rskip BWT for the out side, and we support the occurrence operation via our standard linear search of the directory for the in side, as when converting converting `(syncmer,offset)` pairs to raw `sym`s. During building this means that we need separate counts for incoming and outgoing edges for each side, but after inserting paths in both directions these become equal, and we only keep a single value in the static representation.

### 3.3 Storage in ONEcode files

The syng graph structure is stored in a .1gbwt ONEcode file. Drawing on 30 years of bioinformatics experience, ONEcode provides a flexible, efficient framework for bioinformatics data storage, implemented with Gene Myers at github.com/thegenemyers/ONEcode and used for example by PathPhynder[12] and FastGA[14]. It supports strongly-typed record-based binary files with built-in schemas, indices and data-specific compression (separate codec adaptively trained per record type), and a simple dependency-free interface (single C source file and header), thread support and standard text interconvertibility. The `gbwt` ONEcode schema support serialisation of the vertex, edge and path structure stored in syng vertex and Rskip structures, effectively equivalent to the information stored in a GFA file[1]. Although syng operates on syncmer graphs, a .1gbwt file can store an arbitrary bidirectional sequence graph with paths over it. In parallel, syng uses a .1khash file type to store the syncmer sequences, enabling fast reconstruction of its syncmer data structures.

## 4 Results

I demonstrate the performance of the Rskip data structure using 92 human genomes comprising the Human Pangenome Reference Consortium (HPRC) release 1[10] minus the two HG002 genomes which will be used as test sequences for alignment. These contain 37,269 sequences totalling 277.4 Gbp, which generated 234 billion instances of 193 million syncmers, with average coverage 120.8, in 37 minutes starting from FASTA sequences. The syncmer hash table and index take 10.4 GB in memory, but can be stored in 4.0 GB on disk as a .1khash file.

It took syng 52 minutes on a Linux server, single-threaded, to build a full bi-directional GBWT from the syncmer lists, using 15.7 GB max memory. Time to add genomes increased from ~ 22 seconds per genome to ~ 40 seconds per genome (Figure 3). The resulting graph contained 339.8 million simple vertex sides connected to only one edge, and 46.2 million with multiple edges that required Rskip objects. Of these 46 million were stored as `Linear` arrays with on average 2.4 symbols and 5.4 runs taking 2.0 GB memory, and 175 thousand as `Dynamic` arrays with on average 53.5 symbols and 410 runs taking 7.8 GB (9.8 GB total). The total BWT length (sum of runs) was 4.2 billion for `Linear` nodes and 1.1 billion for `Dynamic` nodes, with an average run length of 16.4 across both types. In total there were 229 million edges, but the incidence distribution has a very long tail due to complex vertices from repetitive regions such as centromeres: the vertex side with the most edges had 13,506 and the maximum number of runs for one edge was 191,212 with maximum total count 1,274,525. There were 151,991 sides with more than 20 edges. The expected directory list search length was 6.3. On disk, the resulting .1gbwt file used 5.8 GB.

**Fig. 3.**
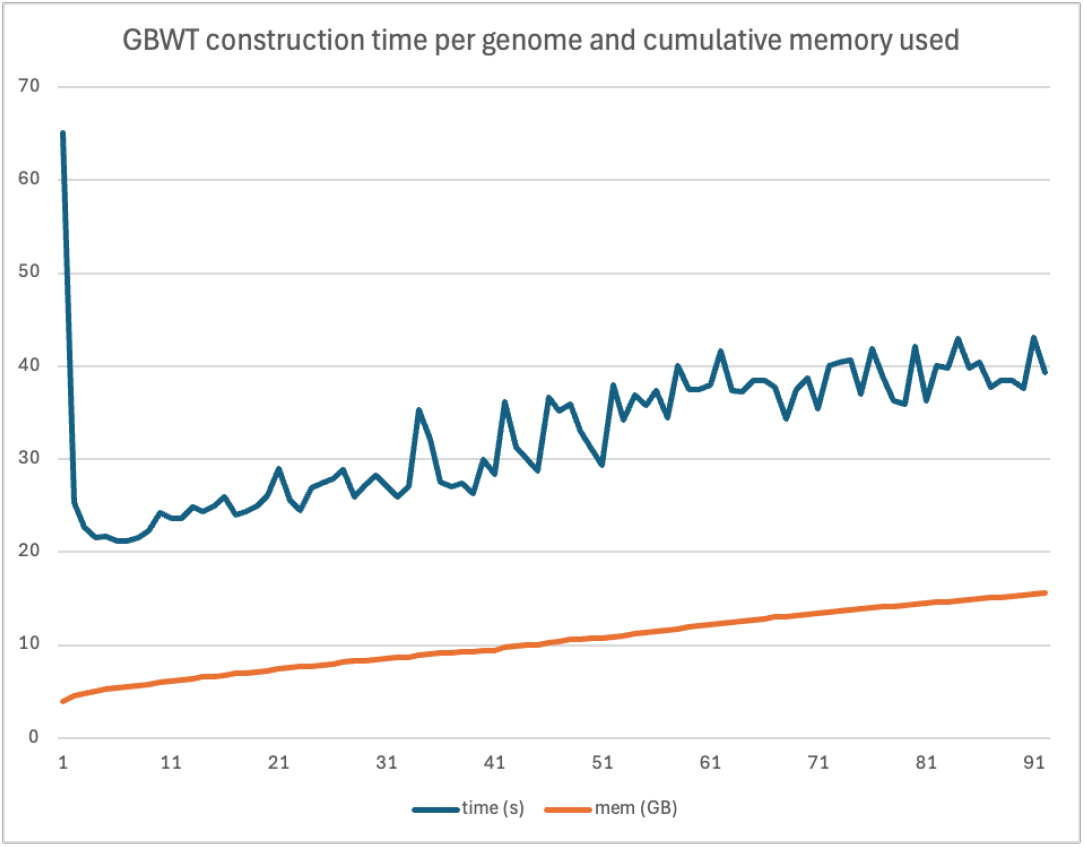
Construction time per genome and cumulative memory used when building a syng pangenome from 92 HPRCv1 genome sequences.

For initial mapping experiments I used the Pacific Biosciences HiFi readset from individual HG002 available from s3-us-west-2.amazonaws.com/human-pangenomics/T2T/HG002/assemblies/polishing/HG002/v1.1/mapping/ hifi_revio_pbmay24/hg002v1.1_hifi_revio_pbmay24.bam, comprising 12.8 million reads totalling 205Gbp (average length 16.0kb). In static mode for searching, the Rskip data structures takes less memory, 1.4 GB for `Linear` and 2.6 GB for `Fixed` (4.0 GB total), both because there is no free space for expansion and because `Fixed` Rskip nodes are smaller than `Dynamic` nodes. With eight threads it takes 14.0 seconds to load the syncmer table and 36.0 seconds to load the vertices, edges and GBWT data. A single forward pass search of the 205 Gbp took 468 seconds or 2.3 seconds per Gbp, coming to 8.6 minutes total for the search including loading tables. Parallelisation performance did not scale with thread number, presumably because of cache contention accessing both the syncmer and GBWT tables: single-threaded times were 125.6 seconds to load the graph (3.5x) and 1890 seconds to search (9.2 seconds per Gb, 4x).

This scan found 204 million maximal exact matches (MEMs) of average length 1304bp, with only 249 reads failing to find matches. MEMs are terminated either by a syncmer not found in the pangenome reference, which may be caused either by a sequencing error or a true genetic variant, or by ancestral recombination generating a new haplotype not present in any reference sequence. In this case I believe the primary reason is sequencing error. We know that the PacBio HiFi sequencing error rate is around 0.1%, and that most errors are in homopolymer run lengths. When I homopolymer compress both the references and the reads, resulting in a 30% length reduction, then average match length increases to 4421bp compressed, equivalent to approximately 6300bp in original bases.

Finally, I examined the empirical length distribution of the linear search step during the `rank()` operation during MEM finding for the HG002 haplotype 1 human genome implemented over the `Dynamic` Rskip structure. In this case, 89.5% of `rank()` operations on `Dynamic` nodes did not require a linear search: the `rank()` operations was already located at the correct symbol. For the other 10.5%, the mean number of search steps taken varies with node side, but on average scales as approximately 0.45*S* where *S* is the number of symbols, or equivalently the number of edges incident to that node side (Figure 4). Across all 34 million `rank()` calls for this genome scan on skip list GBWTs, i.e. those more than 128 runs long, the average linear search count was 25.6.

**Fig. 4.**
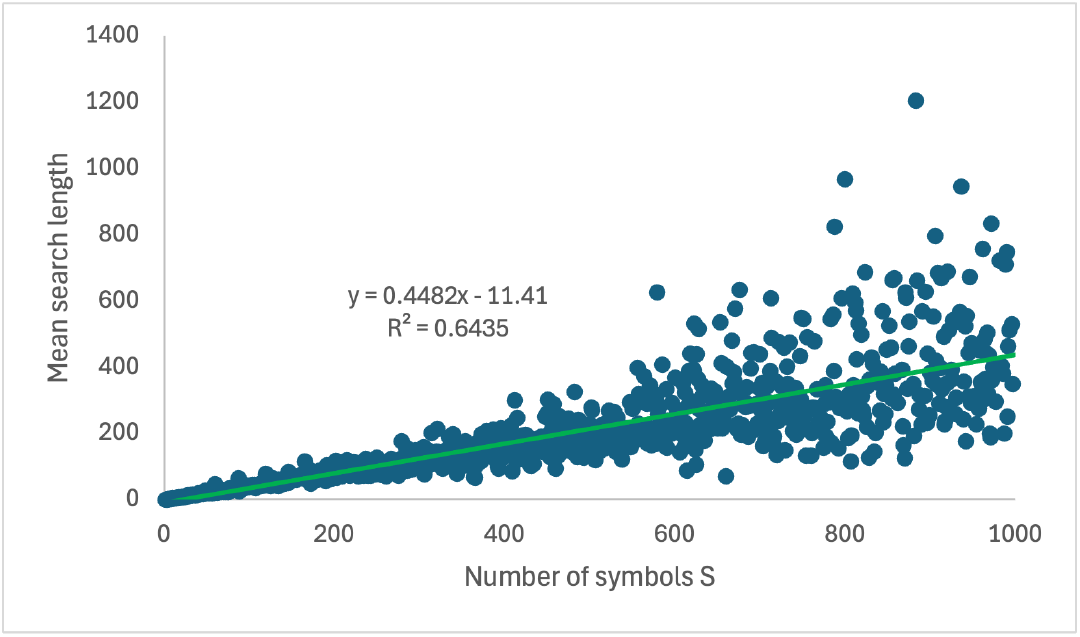
Dependency of the number of linear search steps during general `rank()` on the number of symbols *S*. A point is plotted for each value of *S* from 2 to 1000, at the mean across all `rank()` calls for which *S* has that value, while mapping HG002 haplotype 1 to the HPRCv1 pangenome.

## 5 Discussion

I have presented a GBWT computational model and implementation built on 𝒪 (log *R*) core operations, which operates efficiently at the scale of a hundred human genomes. Empirical time complexity growth as seen in Figure 3 depends on many factors beyond run lengths, but is increasing relatively slowly and apparently sublinearly, so it is reasonable to expect this implementation to scale to the thousand haplotype level envisioned in current pangenome projects.

Notably, `syng` construction is much faster than the time taken by Minigraph-Cactus or PGGB to construct the HPRCv1 `vg` graphs[10]. For example Minigraph-Cactus took approximately 3 days using 25 32-core 256 GB servers (except for one step requiring 512 GB servers) [8], while PGGB took 18.5 hours and 138 GB using 48 threads to build just chromosome 6, less than 1*/*17 of the whole human genome[7]. That said, the `syng` graphs are fundamentally different in that the Minigraph-Cactus and pggb graphs are based on multiple sequence alignments that aim to identify large scale colinear chromosomal homology with few if any cycles and to represent variants at the single base level, whereas the syncmer graph built by `syng` contains many cycles through repetitive sequences and works at a resolution of the syncmer length.

Similarly the GBWT described here is fundamentally different to the one implemented in the `vg` package that supports search using `giraffe`[23]. The DNA sequences of the vertices in `vg` graphs are not unique, and so the `giraffe` GBWT must index the `{A,C,G,T}` strings, rather than sequences of graph vertices, which in our case are syncmers each with their own distinct sequence so indexable by sequence.

Our construction is much closer to a de Bruijn graph as used in sequence assemblers[16, 9], but only using a covering set of sparse *k*-mers as in Ye et al.’s sparse de Bruijn graph assembler[25], MBG[18] and Verkko[19]. Indeed, the syncmer code in `syng` (though not `rskip`) has already been used in the `syncasm` assembler that forms part of the Organelle Assembly ToolKit `oatk`[26]. Another assembler, `mdBG`[5], builds a de Bruijn graph over minimizers and explicitly discusses the application of building pangenomes from bacterial genome sequences. However, `mdBG` differs from `syng` and other assemblers mentioned above in not using overlapping *k*-mers and so not representing all the sequence in the graph. Further, none of these assemblers store the complete paths of all the input sequences in succinct form, as our `rskip` data structure does.

The MEM finding results presented above are only an initial step towards sequence searching. However, enough matches were found to each read to act as seeds for more complete alignment that accepts sequencing errors and mutations, perhaps using principles from Myers’ wave alignment algorithm[13].

In the long run, as indicated in the introduction, the goal is to support imputation of genome sequences from partial data such as low coverage or short read data sets together with a pangenome reference panel. This will require pulling out the most likely pair of haplotype sequences through the graph, based on coverage of read alignments together with the longer range haplotype structure represented by the paths stored in the GBWT.

## Acknowledgments

I thank Chenxi Zhou and Gene Myers for ideas and comments relating to this work, and more generally many other colleagues over the years for stimulating relevant discussions. This work was supported by Wellcome Discovery award 317408/Z/24/Z and a grant on Quantum Pangenomics from Wellcome Leap’s Q4Bio program.

